# Non-Canonical Role of DNA Mismatch Repair on Sensory Processing in Mice

**DOI:** 10.1101/2025.02.13.638164

**Authors:** Sadia N. Rahman, Demetrios Neophytou, Siboney Oviedo-Gray, Bao Q. Vuong, Hysell V. Oviedo

## Abstract

DNA repair mechanisms are essential for cellular development and function. This is particularly true in post-mitotic neurons, where deficiencies in DNA damage response proteins can result in severe neurodegenerative and neurodevelopmental disorders. One highly conserved factor involved in DNA repair is Mut-S Homolog 2 (*Msh2*), which is responsible for correcting base-base mismatches and insertion/deletion loops during cell proliferation. However, its role in mature neuronal function remains poorly understood. This study investigates the impact of *Msh2* loss on sensory processing in a mouse model. Using electrophysiological and molecular assays, we identified significant deficits in cortical and thalamic sound processing in *Msh2^-/-^* mice. These deficits were linked to dysfunction of the thalamic reticular nucleus (TRN), a brain region that critically regulates corticothalamic and thalamocortical activity. Our findings revealed increased oxidative damage, aberrant neuronal activity, and elevated parvalbumin (PV) expression in PV^+^ interneurons in the TRN of *Msh2^-/-^* mice. Additionally, we observed the presence of connexin plaques, indicating that disrupted gap junction formation may contribute to impaired TRN function. These results underscore the critical role of *Msh2* in supporting the functionality of PV^+^ interneurons in the TRN, thereby profoundly influencing sensory processing pathways. This study provides new insights into the importance of DNA repair mechanisms in neuronal development and function, potentially contributing to our understanding of their role in neurological disorders.

## Introduction

A critical function of cells is their ability to recognize and repair genotoxicity over the course of a cell’s lifetime, as it may accumulate close to 10^5^ lesions per day^1^. This is especially crucial in the context of post-mitotic cells, such as neurons. In these cells, the abrogation of different proteins involved in the DNA damage response often leads to a wide range of severe neurodegenerative and neurodevelopmental disorders, such as xeroderma pigmentosum, Cockayne syndrome, and ataxia telangiectasia^2^. The critical role of DNA repair pathways in maintaining genome integrity and their importance in the development and maintenance of the nervous system underscore their significance in preventing neurological disorders^2,3^. One highly conserved repair pathway, mismatch repair (MMR), is used by cells to target base-base mismatches and insertion-deletion loops (IDLs)^4^. Monoallelic deficiencies in any of the MutS homolog proteins (MSH2, MSH3, or MSH6) can result in microsatellite instability, while biallelic mutations in MMR factors (known as constitutional mismatch repair deficiency syndrome) are characterized by agenesis of the corpus callosum (CC), a tendency to form aggressive glioblastomas, and increased oxidized DNA^5–9^. Recent studies also show MMR’s influence in trinucleotide repeat expansion diseases such as Huntington’s disease, myotonic dystrophy, and fragile X syndrome. However, the mechanisms by which MMR contributes to these diseases and syndromes are unknown^10–12^. Moreover, whether or how MMR plays a role in the function of terminally differentiated cells (e.g. post-mitotic neurons) has not been established^13,14^, and this study aims to investigate this question. Unraveling the role of MMR in neural function has the potential to reveal mechanisms underlying neurodegenerative disorders, brain development, and the overall maintenance of neuronal health.

An essential component for the assembly of various recruitment complexes that target sites of mismatch is MSH2 (MutS homolog 2), which heterodimerizes with MSH3 and MSH6 to repair DNA^6^. *In-situ* hybridization data shows that *Msh2* is ubiquitously expressed across different brain regions, including the hippocampus, cortex, thalamus, and hypothalamus^15^.

Broadly expressed genes are more likely to be involved in core functions required for cellular homeostasis, including DNA repair, general transcription, and metabolic regulation^16^. In one of the few studies examining the impact of *Msh2* deficiency on brain function, a notable association was observed with dysmyelination of axonal projections in the corpus callosum of mice^17^.

Nevertheless, MMR is expected to play an important role in glial cells, such as oligodendrocytes, due to their capacity for proliferation, which aligns with MMR’s well-conserved function in DNA replication. However, MMR’s specific involvement and impact on non-proliferative cells, such as neurons, are not yet well understood.

In this study, we employed a top-down approach to examine the effects of *Msh2* deficiency on neuronal function. Deficits arising from neurogenetic disorders can present as weak or noisy processing deficits in ascending neural pathways, gradually accumulating and leading to significant high-level dysfunction^18^. We screened for auditory processing deficits in adult *Msh2^-/-^* mice^19^ using a combination of electrophysiological and molecular assays. We observed widespread decrease in sound-evoked activity in the auditory cortex (ACx) of *Msh2*^-/-^ mice, and hypothesized dysfunction in subcortical regions that play a critical role in modulating cortical and thalamic activity. Specifically, we implicate thalamic reticular nucleus (TRN) pathology as a possible cause of sensory processing deficits in *Msh2^-/-^* mice. Molecular assays revealed that abnormal cellular function and oxidative damage were significantly enhanced in the TRN, and electrophysiological recordings uncovered aberrant patterns of neural activity.

Altogether, these data suggest that *Msh2*, and therefore MMR, play an important role in the development or maintenance of the TRN, thereby supporting downstream cortical processing.

## Methods

*Msh2-*deficient mice (*Msh2^-/-^*) were a gift from H. te Riele. All mice were at least 8 weeks of age. All procedures follow vertebrate animal protocols approved by CCNY IACUC.

### Auditory Stimulation

Mice were placed individually into a sound-attenuated booth. All mice were provided with water, food, and air circulation. The mice remained in the booth for at least 5 hours before being presented with a free-field stimulation (Avisoft Bioacoustic speakers, Glienicke Germany) of frequency modulated (FM) up and down sweeps of 1-40 kHz at 2 octaves/sec. The stimuli were 2 seconds long and separated by an interval of 18 seconds of silence between the end of one stimulus and the beginning of another. This procedure continued for 30 minutes in total. After stimulation, the mice were kept in the sound booth for an additional 50 minutes post-stimulation prior to perfusion.

### Preparation of Brain Sections

Mice were given an intraperitoneal (IP) injection of ketamine (75mg/kg) and medetomidine (0.5mg/kg) in 0.9% NaCl. After anesthesia was confirmed by the absence of a reflexive twitch, mice were perfused intracardially with 15mL of ice-cold phosphate buffered saline (PBS, 1.37 M NaCl, 27 mM KCl, 100mM Na_2_HPO_4_, 18mM KH_2_PO_4_). Mice were then perfused with 20mL of 4% paraformaldehyde (PFA, Alfa Aesar). Post perfusion, mice were decapitated and dissected brains were placed in 4% PFA to post-fix for 48 hours. 50µm free floating sections were prepared on a vibratome on a horizontal or coronal plane.

### Immunohistochemistry

Sections were washed with PBS (3x for 15 minutes) at room temperature (RT) and gently rocked for each wash before being blocked (5% goat serum, 0.3% TritonX-100 in PBS) for 2 hours at RT. Following blocking, sections were incubated in primary antibodies (anti-parvalbumin, Sigma-Aldrich, P3088, 1:500; anti-parvalbumin, Cell Signaling, E8N2U, 1:1,000; anti-cfos, Synaptic Systems, 226-003, 1:2,000; anti-cx36, ThermoFisher Scientific, 37-4600, 1:250; anti-acrolein, Abcam, AB240918, 1:200 at 4°C overnight. Sections were then washed with PBS (4x for 15 minutes) and then incubated with secondary antibodies for 2 hours at RT (Alexa-Fluor 546, Alexa-Fluor 488). Unbound secondary antibody was washed in PBS (3x for 15 minutes) at RT. A subset of sections were then counterstained with 4’, 6-diamidino-2-phenylindole (DAPI) in PBS (1 µg/ml) for 15 minutes after the final wash of PBS. Sections were then mounted using Fluoromount-G (eBioscience) and imaged through confocal microscopy (Zeiss LSM 800/Zeiss LSM 880).

### Cellular Quantification/Analysis

Quantification of immunohistochemistry was conducted utilizing Fiji/ImageJ. All images were subject to consistent parameters during the imaging process, such as the same exposure time, Z-stack intervals, and post-processing/editing in Fiji. Quantification of cfos*-*positive cells was performed by selecting a region of interest, cfos-positive cells were then counted using the Cell Counter plugin available on ImageJ/Fiji. Quantification of PV-positive (PV^+^) cells was conducted by choosing 3 representative regions of interest (ROI) and manually counting all PV^+^ cells in that region. Cell count was then divided by the area (mm^2^) and averaged over 3 regions. Quantification for cellular fluorescence was conducted by selection of 50 representative cells.

Background fluorescence was subtracted from measured integrated density and averaged over 50 cells. Quantification for PV^+^-specific connexin staining was conducted with CellProfiler, by creating a mask of PV-cells and only measuring puncta fluorescence in aforementioned masked PV^+^-cells, removing background fluorescence by using non-PV^+^ cell tissue as a baseline, running an Otsu three-class threshold, and normalizing by area of the cell. Quantification of acrolein fluorescence in the TRN was also conducted with CellProfiler. Acrolein-positive cells were readily identifiable in the TRN of *Msh2^-/-^*mice (Figure 7a); therefore, we used the fluorescence signal from the knockout to develop an automated pipeline to identify and quantify acrolein intensity (Otsu two-class threshold and normalizing by the area of the cell). For statistical analysis of image quantifications, a Student’s two-tailed t-test was conducted in GraphPad Prism 9 and Matlab.

### In-vivo Electrophysiology

A total of 2 *Msh2^+/+^* and 2 *Msh2^-/-^* were used for in vivo recordings of the TRN and ACx. Mice were given an IP injection of ketamine (75mg/kg) and medetomidine (0.5mg/kg) in 0.9% NaCl for anesthetized recordings. Anesthesia was supplemented during surgery and throughout the recordings as needed. Following anesthesia, mice were kept on a heating pad at 36-38°C while placed in a stereotaxic instrument containing head-fixed orbital bars, and a bite bar. To target the thalamic reticular nucleus, we made a craniotomy (approx. 3x2 mm^2^) and durotomy centered at 2.07mm lateral to the midline, and 1.3mm posterior to bregma. A 32-channel silicone probe (Cambridge Neurotech) was lowered at a 15° angle relative to the midline, at a depth of 3.2-3.5mm. The probe’s recording sites spanned 775µm. The exposed cortex was kept moist with cortex buffer ((in mM) 125 NaCl, 5 KCl, 10 Glucose, 10 HEPES, 2 CaCl_2_, 2 MgSO_4_) throughout the recording session. To target the auditory cortex, we made a craniotomy (approx. 3x2 mm^2^) and durotomy centered at 1.5mm anterior and 4mm lateral to lambda. The exposed cortex was kept moist with cortex buffer. The probe was inserted into the auditory cortex at a depth of 800 µm±100 µm from the tip of the probe. The probe’s recording sites spanned all layers of the cortex.

Recordings were obtained using Cheetah software (Neuralynx), with all data sampled at 31kHz. All recordings were done in a sound-attenuated chamber, using a custom-built real-time Linux system (200kHz sampling rate) driving a Lynx-22 audio card (Lynx Studio Technology, Newport Beach, California, USA) with an ED1 electrostatic speaker (Tucker-Davis Technologies, Alachua, Florida, USA) in a free-field configuration (speaker located 6 inches lateral to, and facing the contralateral ear). Recordings from the thalamic reticular nucleus were done in the absence of stimuli to assess spontaneous activity. Cortical recordings were stimulus driven. The stimuli were created with custom MATLAB scripts to compute tuning curves. We used a stimulus set that contained: a suite of pure tones (16 frequencies, 3 amplitudes: 20, 50, 80dB) that lasted 100ms.

### In-vivo Analysis

For analysis of in vivo recordings, we used Kilosort and Phy to sort spikes, extract spike times, and determine spike clusters. Spike clusters were determined to be in the TRN based on their position along the probe relative to the depth that the probe was lowered to. Only narrow spiking neurons were included in the analyses. Following spike sorting and clustering, we used custom made MATLAB scripts to analyze spiking activity, as well as tone responses for cortical recordings. To determine the variability of spike trains in TRN neurons, we calculated the coefficient of variation of inter-spike intervals (ISIs), formally known as the Cv2 metric. The calculation for Cv2 was first described in Holt et al^20^, and is as follows: (2 × |Δt_(n+1)_ – Δt_n_ |)/|Δt_(n+1)_ + Δt_n_ |, where Δt_n_ describes the ISI. To analyze cortical recordings, we computed tuning curves for all clusters. After computing tuning curves, we determined their onset firing rate by counting the number of spikes within a 100ms period after the start of the stimulus. We calculated tone responsive fields (TRFs) to the tones using these firing rates. Intensity thresholds were then determined by measuring the lowest intensity at which each cluster responded with a time-locked response in 25% of the trials presented at that intensity (as previously described in South & Weinberger^21^). Bandwidths were also determined for each cluster at 50dB and 80dB by using the lowest and highest frequency at which the clusters responded in 25% of the trials presented. In general, we observed fewer tone responsive clusters of neurons in *Msh2^-/-^* compared to *Msh2^+/+^* across the same number of recording sites. Therefore, for all analyses we randomly sampled clusters from *Msh2^+/+^* mice to match the same number of clusters analyzed in *Msh2^-/-^* animals. For these analyses, cluster outliers were removed after matching the number of clusters across genotypes.

## Results

### Thalamo-cortical loss of sound processing function in Msh2-null mice

The brain’s capacity to process sensory information is vital, relying on the precise and efficient interaction of cellular and circuitry mechanisms. Considering the potential role of MMR in neuronal function and development, we postulated that the loss of *Msh2* would initiate a cascade of neuronal dysfunction, which would be especially pronounced in higher-order sensory regions like the auditory cortex^18^. Changes in sensory input cause an increase in neuronal activity, which can lead to elevated expression of immediate early genes. Therefore, we stained for the protein product of the immediate early gene *cfos* to detect recently active neurons. To screen for changes in sound-evoked responses as a result of *Msh2* loss, we presented auditory stimuli (Figure 1a) to evoke neuronal activity in the ACx and quantified cfos^+^ neurons. Using frequency sweeps, we stimulated *Msh2^+/+^* mice and observed the previously reported lateralized activity, indicated by the increased number of cfos^+^ cells in layers 2/3 (L2/3) of the right ACx compared to the left ACx^22^ (Figure 1b (i, ii)). In the absence of *Msh2*, however, a significant decrease in cfos^+^ neurons was observed in both auditory cortices, and lateralized activation of the auditory cortices was also abolished (Figures 1b (iii, iv) and 1c). These results suggest that the loss of *Msh2* leads to significant hypoactivity and impairment in normal sensory processing functions within the ACx.

**Figure 1:**
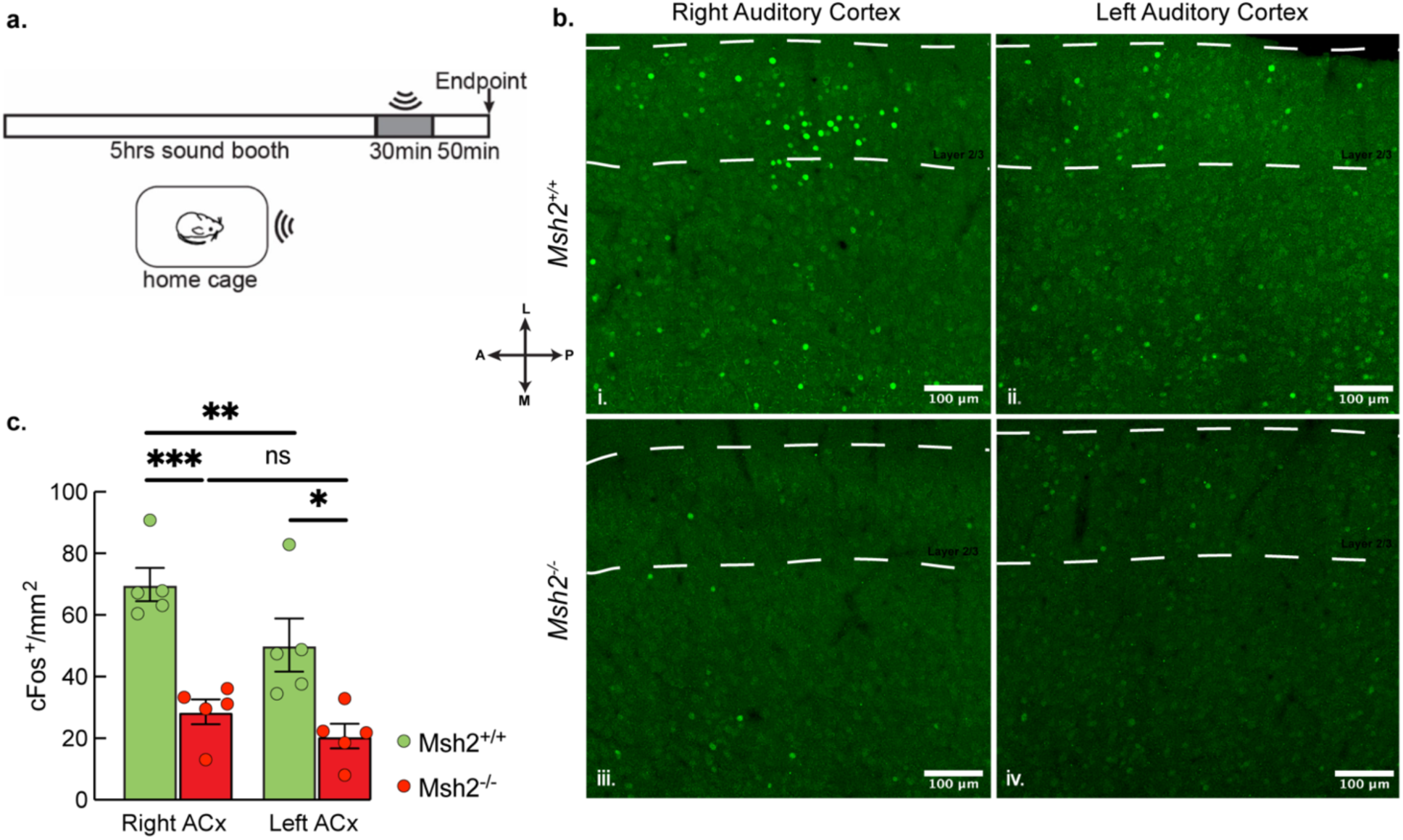
Diminished auditory function in the ACx of *Msh2^-/-^* mice. **a)** Schematic of sensory function assay utilized: mice were placed in a sound-proof chamber for 5 hours to acclimate, given 30 minutes of auditory stimulation, and then remained in the cage for 50 minutes post-stimulation prior to perfusion. **b)** cfos expression in the right and left ACx in *Msh2^+/+^*(i, ii), and *Msh2^-/-^* (iii, iv) mice. Dotted lines indicate layers II/III of the cortex. Scale bars are 100µm. **c)** Population data showing that there are significantly less cfos^+^ neurons in the right and left ACx of *Msh2^-/-^* compared to *Msh2^+/+^*littermates. Statistical analysis was conducted through a Student’s two-tailed unpaired t-test (*, p < 0.05; **, p ≤ 0.001; ***, p ≤ 0.0001; n = 5 mice for each genotype, error bars are SEM). Stereotaxic axis shows anterior-posterior (A/P) and medial-lateral (M/L) axis.

To examine in greater detail the extent of sound processing deficits in the ACx of *Msh2^-/-^* mice, we recorded multiunit neural activity using silicon probes that spanned L2-L6 (Figure 2a). Multiunit responses were sorted using Kilosort (see Methods) to identify clusters of spikes from putative single units. Interestingly, we observed tone-evoked firing in *Msh2^-/-^* animals, albeit significantly lower than in *Msh2^+/+^* (Figures 2b and 2c). We examined differences in frequency encoding between the genotypes by analyzing the tone-responsive fields of single units. We found a significant disparity in intensity thresholds between the two genotypes (Figure 2d). This finding indicates a considerable increase in the sound level intensity required to elicit sound-evoked responses in *Msh2^-/-^* mice. In addition, we examined potential differences in the frequency response bandwidth. This metric captures the span of frequencies over which there is a measurable response above threshold. We performed this analysis at 50dB and 80dB (termed BW30 and BW60) in clusters of both genotypes (Figure 2e). While there were no significant differences at BW60, the BW30 was significantly narrower in *Msh2^-/-^* (1.48 octaves) compared to *Msh2^+/+^* clusters (2.29 octaves). Together, these results suggest a reduced dynamic range in encoding frequency information, compromising the network’s ability to respond to simple and complex sounds. The cfos data and direct recordings of neural activity reveal significant and widespread deficits in sensory-evoked activity in the ACx of *Msh2^-/-^* mice and point to possible dysfunction in subcortical processing stations.

**Figure 2:**
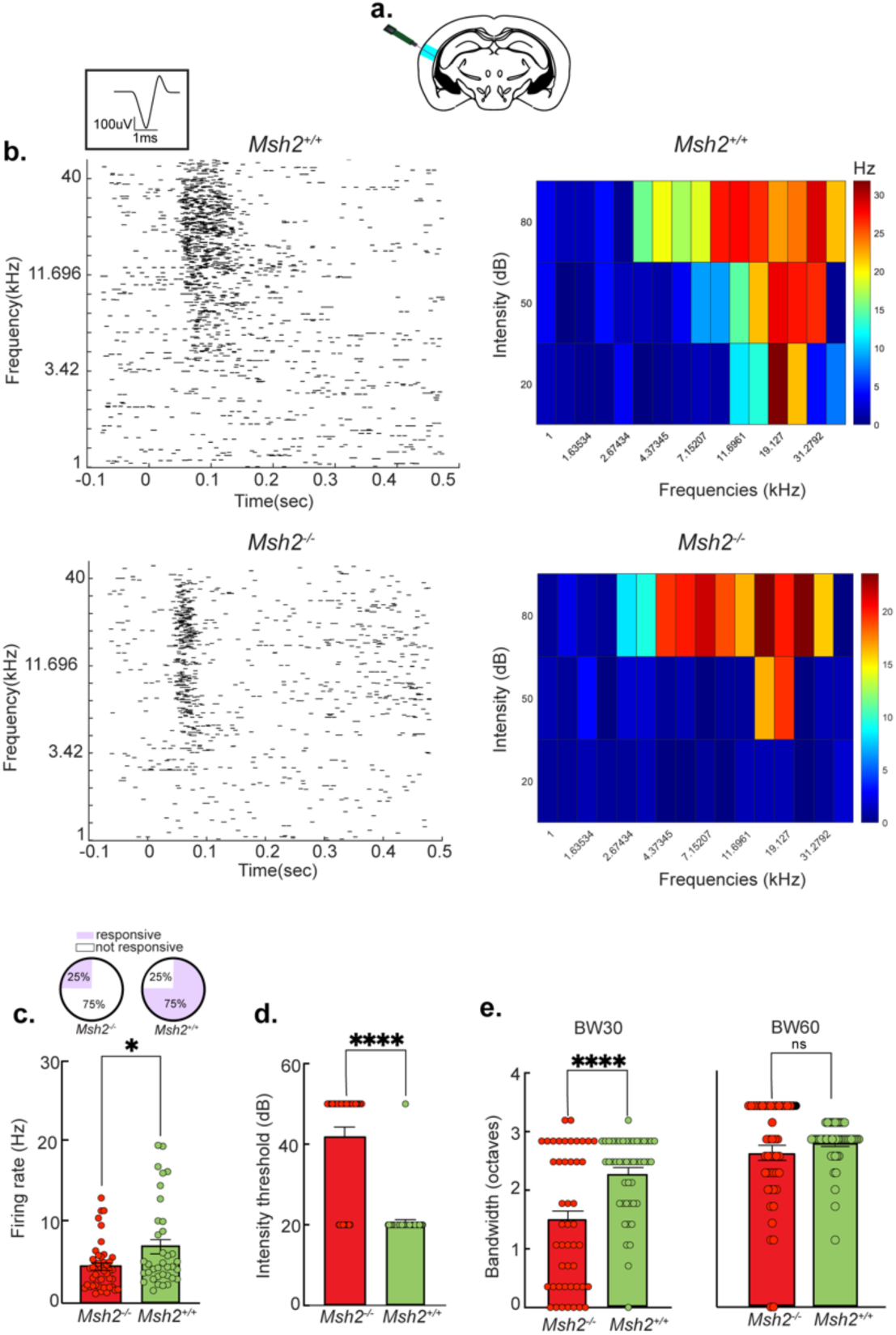
Deficits in frequency response properties in the ACx of *Msh2^-/-^* mice. **a)** Schematic of experimental paradigm to record from the ACx (shown in cyan) with multi-channel silicone probes. **b)** Representative examples of rasters (left) and tone responsive fields (TRFs; right) from individual clusters of *Msh2^+/+^* (top) and *Msh2^-/-^*(bottom) mice. Inset is a representative spike waveform from a putative regular spiking neuron in the ACx. **c)** Proportion of tone-responsive clusters (top), and average firing rate from responsive clusters in *Msh2^+/+^*and *Msh2^-/-^* mice (bottom, n = 47 clusters for each genotype; *, p = 0.01). **d)** Average intensity thresholds (dB) from *Msh2^+/+^* and *Msh2^-/-^* clusters (n = 47 clusters for each genotype; ****, p ≤ 0.00001). **e)** Average bandwidths (in octaves) of tuning curves at 30dB (left; ****, p ≤ 0.00001) and 50dB (right) above our lowest intensity level (20dB) from *Msh2^+/+^* and *Msh2^-/-^* clusters (n = 36 clusters for *Msh2^+/+^* mice; n = 43 clusters for *Msh2^-/-^* mice). All error bars are standard error of the mean, n = 2 mice for each genotype.

To screen for sound processing deficits in subcortical auditory areas, we examined cfos expression in the ventral division of the geniculate nucleus (MGBv). This analysis was performed in the same animals stimulated with frequency sweeps and quantified for cfos expression in the ACx (Figure 1). The MGBv is an obligatory relay for all ascending auditory information, ultimately projecting to the ACx. We found that *Msh2^-/-^* animals had a significant decrease in cfos^+^ neurons in the MGBv compared to their *Msh2^+/+^* littermates, suggesting decreased neural activity in ascending auditory pathways (Figure 3). Interestingly, the number of cfos^+^ neurons in the inferior colliculus (IC), an area that provides auditory input to the MGBv^23^, was not significantly different between *Msh2^-/-^*and *Msh2^+/+^* animals (Supplementary Figure 1). Based on this evidence, we reasoned that the sensory processing deficits we see in *Msh2^-/-^* ACx could arise from intra-thalamic processing.

**Figure 3:**
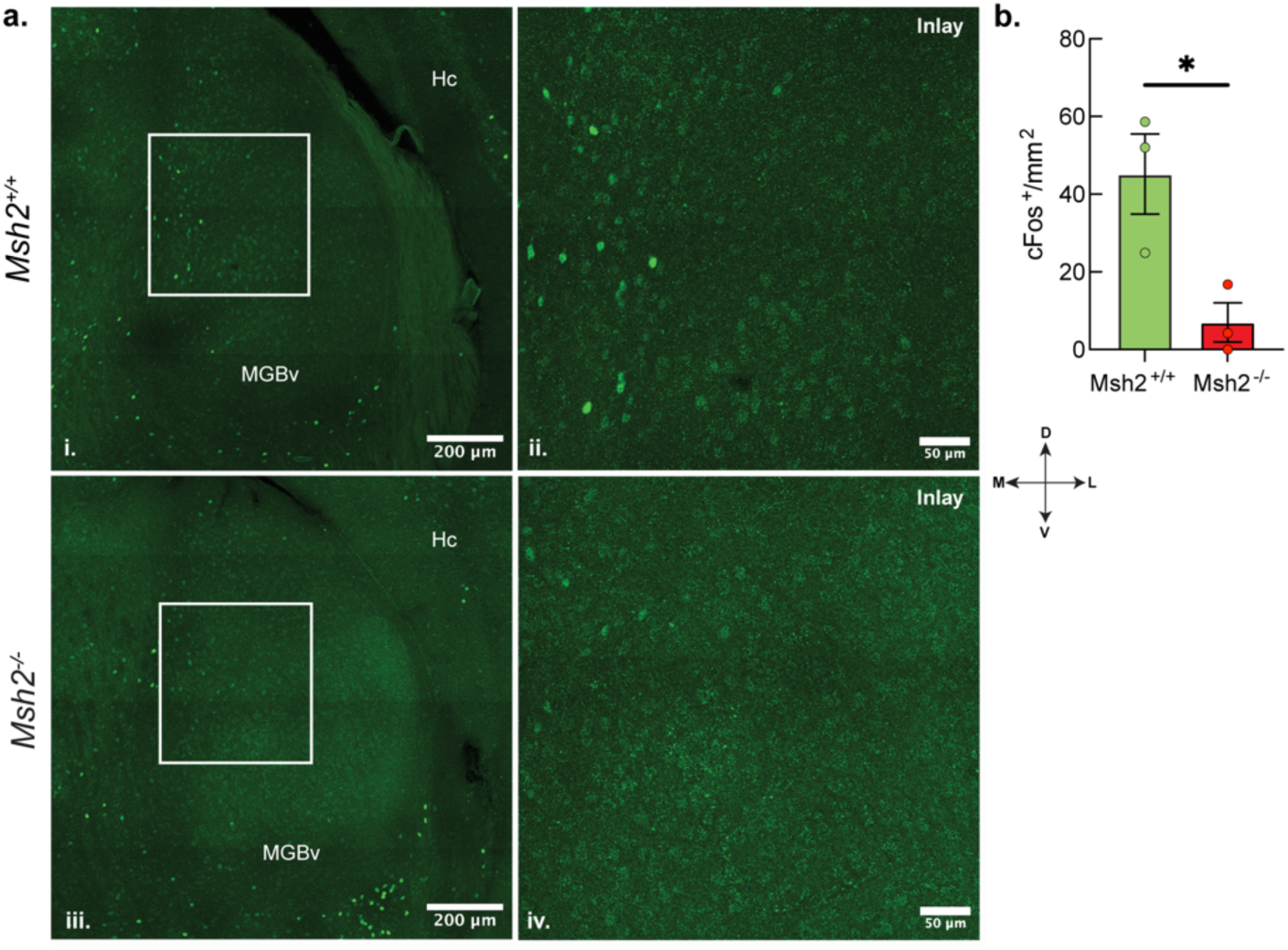
Reduced neural activity in the MGBv of *Msh2^-/-^* mice. **a)** Quantification of cfos^+^ neurons in the MGBv of *Msh2^+/+^* (top, i-ii) and *Msh2^-/-^* (bottom, iii-iv) mice. Scale bars indicate 200µm and 50µm for the inlay. **b)** Statistical analysis of MGBv cfos data conducted through a Student’s two-tailed unpaired t-test (n = 3 mice for each genotype; *, p < 0.05). Stereotaxic axis represents the medial-lateral (M/L) and dorso-ventral (D/V) axis.

### Thalamic reticular nucleus dysfunction in Msh2^-/-^ mice

We asked whether deficits in thalamocortical (TC) activity could result from dysfunction of the thalamic reticular nucleus. The TRN, a shell of largely GABAergic inhibitory neurons in the dorsal thalamus, gates the bidirectional flow of information between the primary and secondary cortices and first- and higher-order projection neurons^24^. TRN neurons receive glutamatergic inputs from the thalamus and send GABAergic efferents to TC projection neurons, partaking in feedforward and feedback inhibitory circuits involving the sensory cortices and thalamus (Figure 4a).

**Figure 4:**
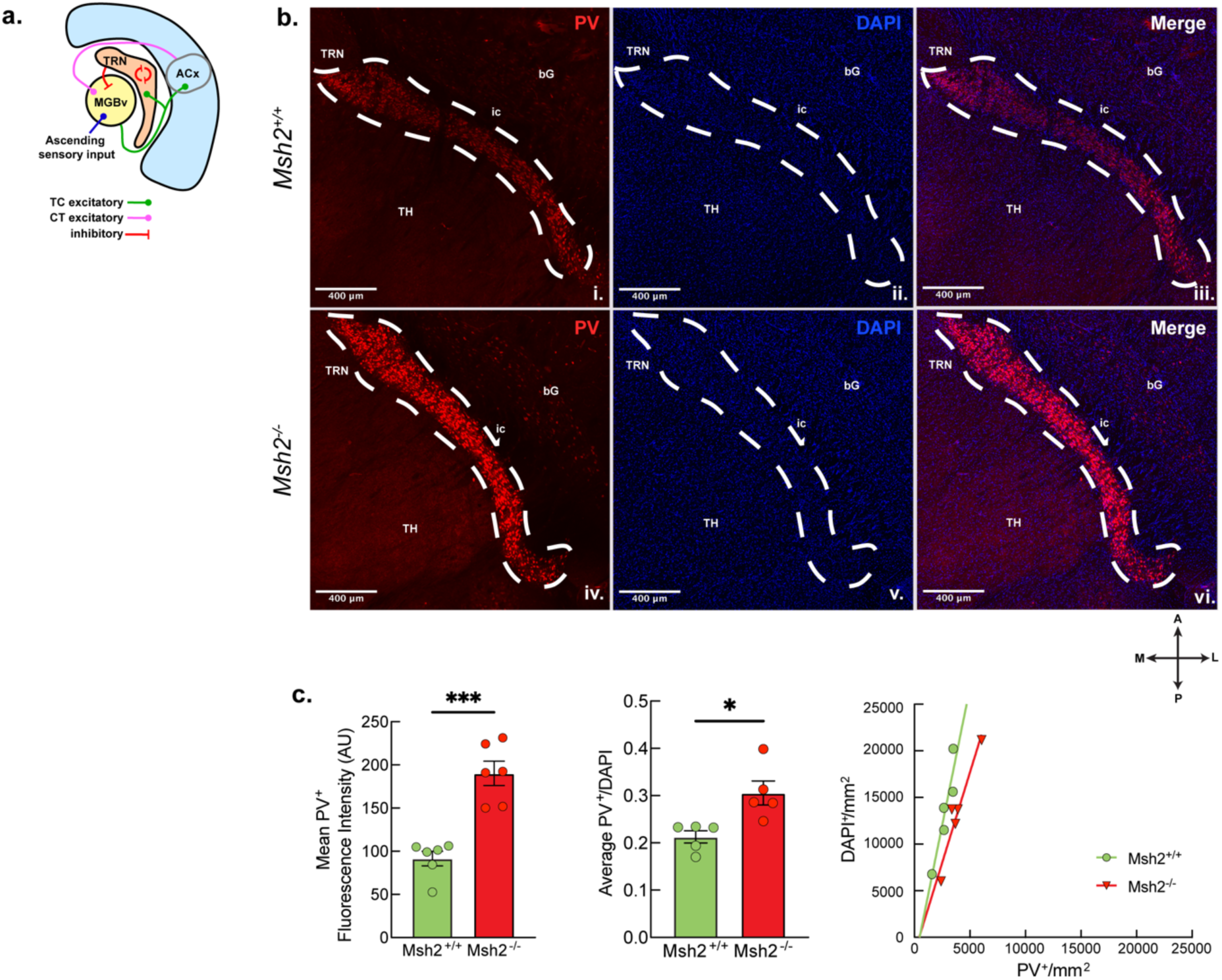
Greater parvalbumin expression within the TRN of *Msh2^-/-^* mice. **a)** Diagram of TRN 0connectivity; excitatory TC connections from the MGBv send collaterals to the TRN and form reciprocal excitatory connections with the cortex. The TRN provides inhibitory signaling to the thalamus and itself. **b)** Parvalbumin (PV^+^) staining counterstained with DAPI in the TRN of *Msh2^+/+^* (top, i-iii) and *Msh2^-/-^* (bottom, iv-vi) mice. Scale bars represent 400 μm. **c**) **Left**: Mean fluorescence intensity in arbitrary units (AU) (***, p = 0.0001, n = 6 mice for each genotype), **middle**: average PV^+^ neurons per DAPI-demarcated cells (*, p < 0.05, n = 5 mice for each genotype), and **right**: relative density of PV^+^ interneurons in *Msh2^+/+^* and *Msh2^-/-^* mice (n = 5 mice for each genotype). Error bars represent SEM. Statistical analysis was conducted through a Student’s unpaired two-tailed t-test. Stereotaxic axis (lower right) represents anterior-posterior (A/P) and medial-lateral (M/L) axis. TH = thalamus, bG = basal ganglia, ic: internal capsule, Hc = hippocampus, TRN = thalamic reticular nucleus, MGBv = medial geniculate nucleus, ventral division.

Most of the neurons in the TRN express parvalbumin, a calcium binding protein necessary for the regulation of calcium dynamics during neuronal activity^25^. Measuring parvalbumin expression serves as a useful metric for identifying plasticity and pathological conditions in the brain^26,27^. Therefore, to screen for TRN deficits in *Msh2^-/-^* mice, we began with an analysis of parvalbumin expression in PV^+^ interneurons. Notably, parvalbumin expression in the TRN of *Msh2^-/-^* mice was significantly increased compared to *Msh2^+/+^* (Figure 4b (i, iv), c (left)). Moreover, quantification of PV^+^ cellular density indicated a significantly higher proportion of PV^+^ neurons compared to total cell count (as indicated by DAPI) in the TRN of *Msh2^-/-^* mice (Figure 4c (middle)). This observation is also supported by the higher number of PV^+^ neurons per mm^2^ of TRN in *Msh2^-/-^*compared to *Msh2^+/+^*, whereas the number of DAPI^+^ cells per mm^2^ of TRN is similar between the two genotypes (Figure 4c (right)). These data indicate that PV^+^ interneurons in the TRN of *Msh2^-/-^* mice are more numerous and exhibit increased parvalbumin expression compared to *Msh2^+/+^* controls. Upon further investigation we also observed that, on average, PV^+^ interneurons in *Msh2^-/-^* mice exhibited a larger cellular area compared to their *Msh2^+/+^* littermates (Figure 5a). This phenotype was specific to the TRN and was not observed in the ACx of *Msh2^-/-^*mice (Figure 5b, e).

**Figure 5:**
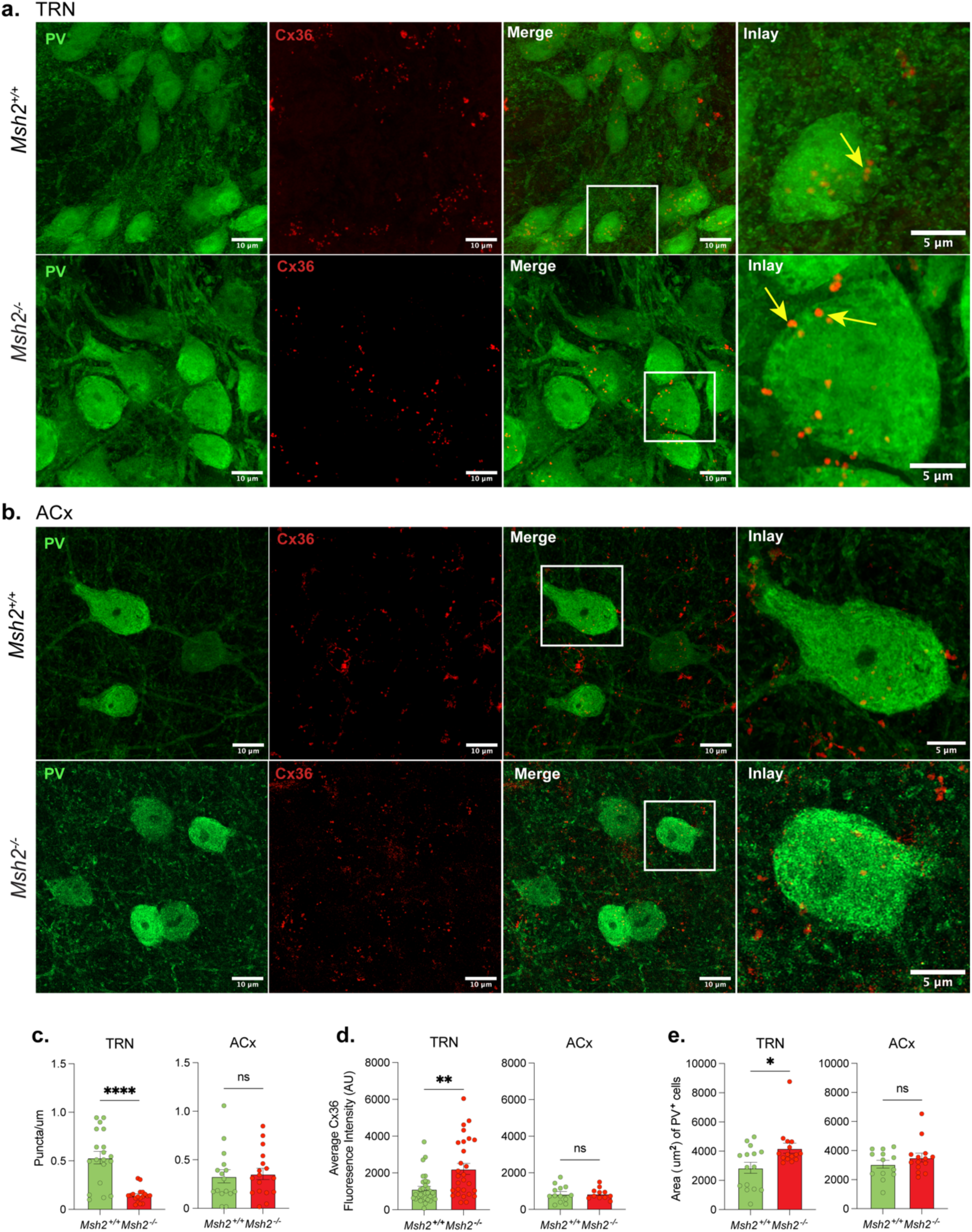
Cellular and molecular abnormalities in PV^+^ TRN neurons of *Msh2^-/-^.* PV^+^ and Cx36 staining in **a)** TRN and **b)** auditory cortices of *Msh2^+/+^* and *Msh2^-/-^*mice. **c)** The formation of Cx36-containing plaques in *Msh2^-/-^*PV^+^ neurons led to a decrease in puncta/µm in the TRN (****, p ≤ 0.0001, n = 3 mice for each genotype; neurons analyzed: 19 from *Msh2^+/+^*and 18 from *Msh2^-/-^*), but not the ACx (neurons analyzed: 16 from *Msh2^+/+^* and 17 from *Msh2^-/-^*). **d)** Cx36 fluorescence intensity (AU) was found to be higher in *Msh2^-/-^*TRN (**, p ≤ 0.01, n = 3 mice for each genotype; neurons analyzed: 30 from *Msh2^+/+^*and 28 from *Msh2^-/-^*), but comparable in the ACx of both genotypes (neurons analyzed: 13 from *Msh2^+/+^* and 11 from *Msh2^-/-^*). **e)** PV^+^ cells in the TRN were also larger in *Msh2^-/-^* animals (*, p ≤ 0.05, n = 3 mice for each genotype; neurons analyzed: 15 from *Msh2^+/+^* and 15 from *Msh2^-/-^*), but not the ACx (neurons analyzed: 13 from *Msh2^+/+^*and 14 from *Msh2^-/-^*). Statistical analysis was conducted through Student’s two-tailed unpaired t-test. All error bars are standard error of the mean (SEM). Scale bars represent 10μm and 5μm (inlay).

Collectively, these abnormal cellular phenotypes prompted us to investigate whether functional features crucial for the TRN’s electrical properties were also affected. This could directly contribute to the disruption of thalamo-cortical sensory evoked responses we observed. PV^+^ interneurons in the TRN are electrically coupled via connexin-36 (Cx36) gap junctions^28^. Therefore, we examined the expression of Cx36 in PV^+^ neurons to determine how the TRN’s role as a modulator of TC activity could be compromised in *Msh2^-/-^*. Like most connexins, Cx36 typically does not remain as a single-unit membrane protein but assembles into connexons-six-protein hemichannels that permit the direct transfer of ions and small molecules between cell bodies^29^. To quantify connexon formation, we measured fluorescence intensity and puncta count per micron of tissue, which reflect the assembly of proteins. In *Msh2^-/-^* TRN, we observed a significant decrease in Cx36 puncta per micron in PV^+^ interneurons (Figure 5c). We also found an increase in Cx36 fluorescence intensity (Figure 5d). This increase was attributed to the formation of aggregates, resulting in Cx36 plaques. These changes in Cx36 were specific to the TRN and were not observed in the ACx of *Msh2^-/-^* mice. The evidence suggests that in *Msh2^-/-^*mice, PV^+^ interneurons in the TRN show altered Cx36 proteodynamics. This disrupted gap junction formation, and consequently interneuron coupling, could compromise the TRN’s modulation of thalamo-cortical activity.

Our observations of high parvalbumin levels and abnormal connexin clustering in *Msh2^-/-^*mice suggest abnormalities in the electrical properties of the TRN, potentially causing the observed downstream dysfunction in auditory processing. To investigate, we recorded spontaneous activity from the TRN *in vivo* to assess possible deficits in intrinsic electrophysiological properties (Figure 6a). The TRN has two characteristic firing patterns: tonic and bursting^30^. Using anesthetized mice, we investigated whether these firing patterns were altered with the loss of *Msh2* by quantifying properties of the interspike interval (ISI). One of the most striking phenotypes in *Msh2^-/-^*was a higher proportion of short ISIs compared to *Msh2^+/+^* (Figure 6b, c). This suggests that in the TRN of *Msh2^-/-^* mice, burst firing epochs become the predominant mode of output, with bursts showing higher spike rates. We also quantified the variability of spike trains using the coefficient of variation of ISIs metric (Cv2). Cv2 can range between 0 and 2, with higher values indicating higher variability in ISIs. Quantifying Cv2 for both *Msh2^+/+^*and *Msh2^-/-^* clusters, we found greater variability in the timing of action potentials in spike trains from *Msh2^-/-^* mice (Figure 6d, e). These findings indicate that the discharge patterns of the TRN, which are essential for generating oscillations and contribute significantly to brain rhythms^31–34^, become disrupted in the absence of MSH2.

**Figure 6:**
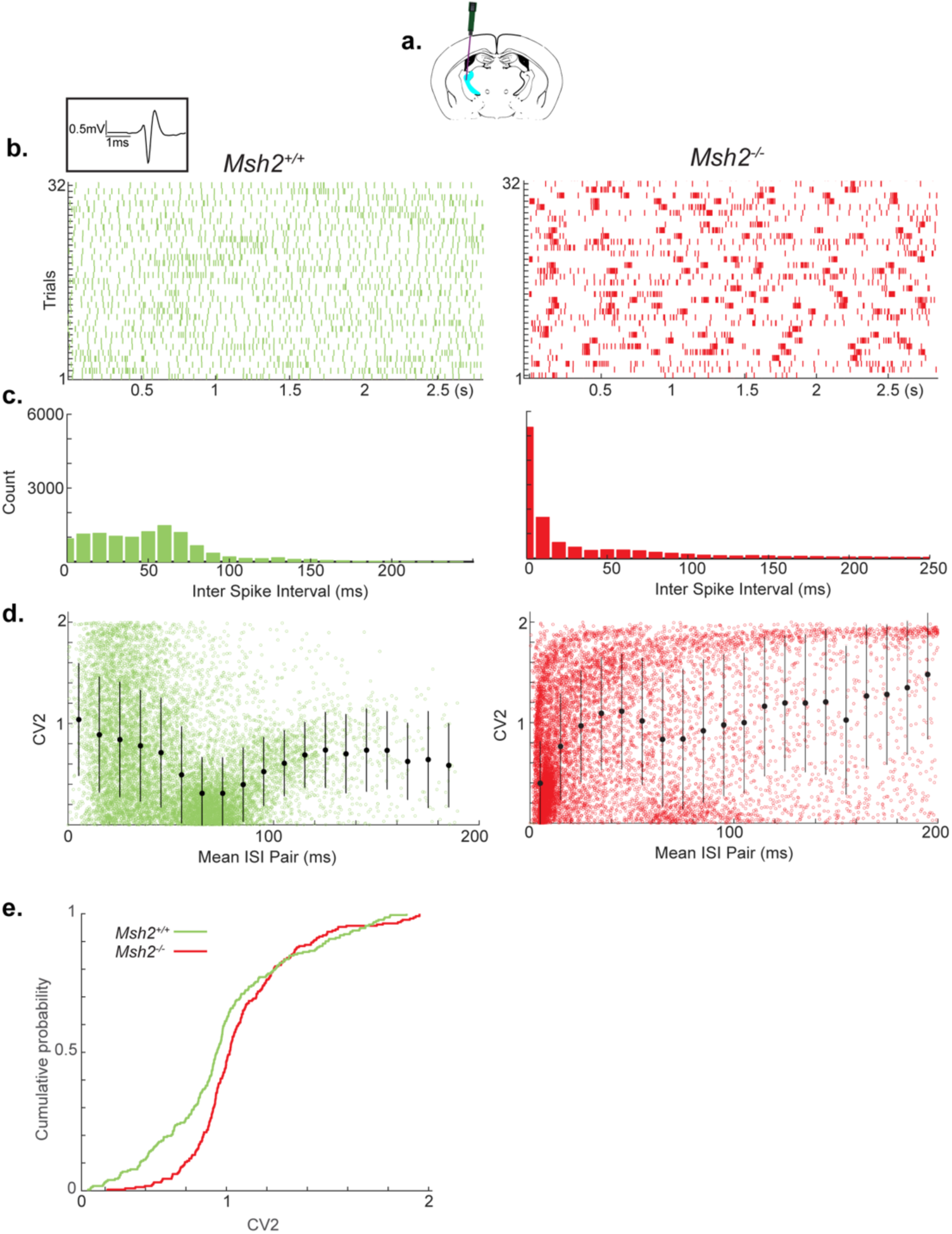
Higher incidence of burst firing in the TRN of *Msh2^-/-^* mice. **a)** Schematic of experimental paradigm to record spontaneous activity from the TRN (shown in cyan) in *Msh2^+/+^*and *Msh2^-/-^* mice. **b)** Representative examples of rasters from *Msh2^+/+^* (left) and *Msh2^-/-^* (right) clusters. Inset is a representative spike waveform from a putative interneuron in the TRN. **c)** Histogram of inter-spike intervals (ISIs) from *Msh2^+/+^* (left) and *Msh2^-/-^* (right) clusters. **d)** Cv_2_ values from *Msh2^+/+^* (left) and *Msh2^-/-^* (right) clusters. Bin sizes are 10ms. **e)** *Msh2^-/-^* clusters have significantly higher Cv2 values than *Msh2^+/+^* clusters (p < 0.05). All error bars are standard error of the mean, n = 2 mice for each genotype.

**Figure 7:**
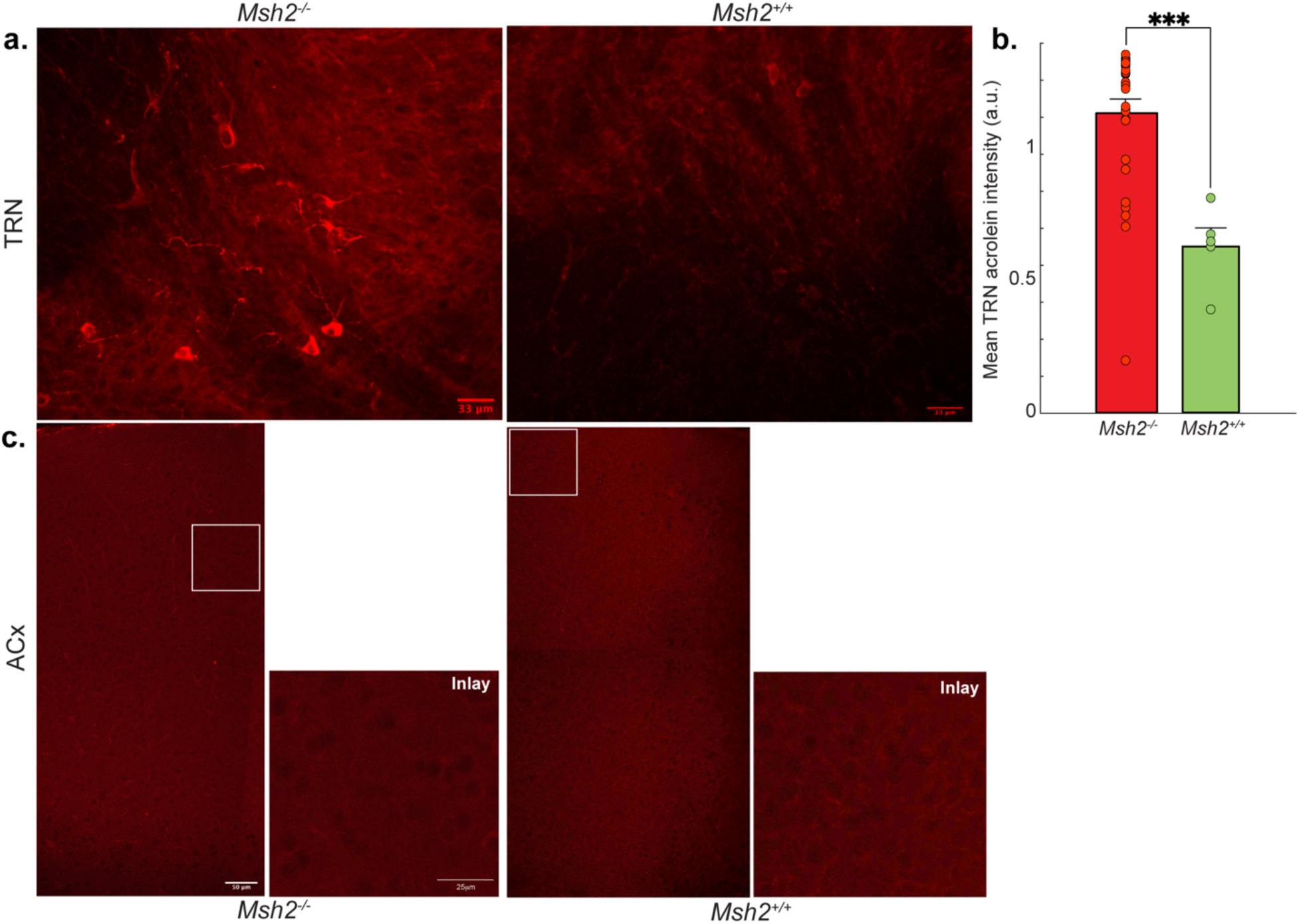
Higher levels of oxidized DNA in the TRN of *Msh2^-/-^* mice. **a**) Staining for acrolein in *Msh2^-/-^* mice (left) revealed a greater number of cells with higher acrolein intensity compared to *Msh2^+/+^* mice (right) in the TRN. Scale bars indicate 33μm. **b**) Mean acrolein fluorescence intensity in the TRN (in arbitrary units, a.u.) is significantly higher in *Msh2^-/-^* mice (*** p = 0.0001, n = 4 mice for each genotype, error bars represent SEM). Statistical analysis was conducted through a Student’s unpaired two-tailed t-test. **c**) Examples of low acrolein levels in the ACx of both *Msh2^-/-^*(left) and *Msh2^+/+^* (right) mice. Scale bars indicate 50μm and 25μm for inlay.

### TRN dysfunction is associated with increased oxidative damage

Collectively, the abnormally high levels of PV, enlarged cell body, connexin plaques, and aberrant electrical activity in the TRN suggest cellular stress caused by the absence of *Msh2*. Because the absence of *Msh2* increases oxidized DNA^9^, we hypothesized that the impaired neural function in *Msh2^-/-^* mice could result from increased reactive oxidative species (ROS). These ROS cause oxidized DNA in TRN neurons, which cannot be repaired. To investigate this, we measured acrolein, a marker of oxidative stress, in both *Msh2^+/+^* and *Msh2^-/-^*mouse brains^35^. Acrolein-positive cells were readily identifiable in the TRN of *Msh2^-/-^*mice but not *Msh2^+/+^* mice (Figure 7a). We implemented threshold parameters (see Methods) to quantify the mean intensity of acrolein-positive cells (Figure 7b). When we applied the *Msh2^-/-^* threshold criteria to the TRN of *Msh2^+/+^* mice, fewer cells passed threshold to be detected as acrolein-positive.

Additionally, the few cells that were detected in *Msh2^+/+^*TRN had significantly lower acrolein intensity compared to the *Msh2^-/-^*TRN (Figure 7b). Cells in the ACx of the *Msh2^-/-^*and *Msh2^+/+^* mice did not pass threshold for acrolein detection (Figure 7c), and therefore did not meet criteria for quantification. Hence, the increased oxidation in the *Msh2^-/-^*TRN, as measured by higher acrolein intensity, shows that ROS levels are elevated in the TRN but not in the ACx of *Msh2^-/-^* mice. This indicates that the loss of *Msh2* may lead to altered TRN function as a result of oxidative stress or oxidative DNA damage.^9,36,37^.

## Discussion

We observed sensory deficits in the absence of MSH2, an essential protein in MMR, using a genetic knockout of *Msh2* in mice. Compared to wildtype littermate controls, *Msh2^-/-^* mice show (1) a significant disruption of sound-evoked neural activity in the ACx and MGBv, (2) elevated parvalbumin expression and PV^+^ cellular density in the TRN, and (3) altered PV^+^ neurons with significantly larger cellular size, abnormal firing properties, and increased Cx36 fluorescence and puncta aggregation, indicating elevated gap junction plaque formation. The abnormal PV^+^ neuronal function correlates with significantly higher levels of acrolein, a marker of oxidative stress, specifically in *Msh2^-/-^* TRN.

In response to frequency modulated sweeps, *Msh2^+/+^* mice showed right ACx lateralized neural activity (indicated by cfos^+^ neurons) in the superficial cortical layers, consistent with findings from previous studies^22,38,39^. In *Msh2^-/-^*, this stimulus-driven response is considerably reduced in the right and left ACx. Using multiunit recordings of neural activity from the ACx, we examined the deficits in stimulus-driven responses in more detail. We found deficits in frequency tuning across multiple metrics in *Msh2^-/-^* mice compared to their wildtype littermates. Indicative of a diminished response to sounds, fewer clusters in *Msh2^-/-^* mice were responsive to tones, and the average firing rate of these responsive clusters was lower, with significantly higher intensity thresholds. Additionally, the dynamic range of their frequency response bandwidth was significantly reduced. To identify the origin of these deficits, we investigated the ascending auditory pathways. Our findings revealed a decrease in sound evoked neural activity in the MGBv but not the inferior colliculus. Based on these findings, we reasoned that the potential sound processing deficits originated within the thalamus. We identified cellular and electrophysiological pathologies in the TRN, which plays a crucial role in modulating thalamocortical pathways and serves as a major source of inhibition for the thalamus. Dysfunction of the TRN has been shown to be linked to neurological disorders with sensory abnormalities, insomnia, and schizophrenia^40^. Despite its importance in sensory information gating, the TRN’s precise cellular and functional organization remain largely unknown.

While we do not discount the possibility of dysfunction in other brain regions resulting from *Msh2* loss, and therefore MMR reduction, our findings pose a fundamental question about the specific vulnerability of the affected PV^+^ interneurons in the TRN. PV^+^ interneuron dysfunction is frequently associated with a multitude of nervous system diseases^26,41–46^. This susceptibility can be attributed to a combination of factors, including genomic instability, high firing rates, and the unique vulnerability of PV^+^ interneurons to a variety of stressors. The TRN is particularly vulnerable, largely because of its predominantly PV^+^ population and its distinction of having some of the highest firing rates in the brain, with firing epochs reaching up to 500 spikes per second^30,47^. As a result, PV^+^ interneurons in the TRN have extraordinary energy requirements to support their firing activity and provide defense against substantial glutamatergic stress. One of the key protective mechanisms employed by these interneurons involves regulating calcium buffering through parvalbumin levels^48^. Our results suggest that interneurons in the TRN appear to be in a high calcium buffering state as evidenced by significantly increased intensity of parvalbumin immunoreactivity (Figure 4c (left)). Moreover, increases in intracellular calcium levels can lead to the induction of DNA damage from reactive oxygen species, which cannot be repaired efficiently in the absence of MSH2^9,36,49^. This exacerbates the impact of genomic instability on the TRN.

We also observed increased gap junction plaque formation in the TRN of *Msh2^-/-^* mice compared to the *Msh2^+/+^* TRN and ACx of both genotypes, suggesting that *Msh2*-deficiency specifically affects the TRN. Connexins are highly dynamic membrane proteins with very short half-lives, lasting only a few hours^50^. The coordination of connexin trafficking and assembly is managed through phosphorylation events controlled by numerous scaffolding proteins and various kinases^50^. Genomic instability can perturb these regulatory mechanisms leading to disruptions in the binding of scaffolding proteins like zona occludens-1 (ZO-1) and an increase in gap junction plaques^51,52^. While these mechanisms have primarily been examined in the context of Cx43 due to its extensive study, the biochemical interactions between ZO and Cx36, the connexin type in the TRN, are well-established^53^. Disruptions in these interactions could, therefore, have similar impacts in the TRN. DNA damage response elements and cellular stress also lead to an increase in the activity of protein kinase C, Akt, MAPK and src kinase, which are all known to phosphorylate connexins and result in an increase in gap junction plaque size and internalization from the plasma membrane^54,55^. Interestingly, an increase in Akt activity could also play a role in the higher density of PV^+^ interneurons we observed in the TRN (Figures 4b and 4c). Akt is a key regulator of neuronal survival, and its overexpression has been shown to prevent apoptosis in postnatal brain development^56^. More relevant to our study, deleting antagonists of Akt activity significantly enhances the survival of postmitotic interneurons derived from the medial ganglionic eminence^57^.

Previous experimental and theoretical studies have elucidated the significance of gap-junctional coupling in mediating burst firing in neural networks^58–61^. An increase in Cx36 gap junction plaques could enhance local coupling between PV^+^ interneurons in the TRN of *Msh2^-/-^* mice. Enhanced electrical coupling has been shown to increase the synchrony of spiking activity in neural networks^62^. Our recordings of spontaneous activity in the TRN revealed more bursting epochs with higher rates of short ISIs in *Msh2^-/-^*mice, consistent with this enhanced electrical coupling (Figure 6). Furthermore, the significantly higher Cv2 values observed in the firing discharge of the *Msh2^-/-^* TRN align with theoretical findings that spike bursts exhibit greater Cv2 values and variability in spike timing^63^. We propose that the abnormal aggregation of Cx36 gap junction plaques, leading to enhanced local coupling, could account for the atypical firing properties observed in the TRN of *Msh2^-/-^* mice. Given that the TRN is an inhibitory nucleus, stronger synchronization of its discharge could more effectively reduce activity in both the MGBv and ACx (Figure 4a)^64^. This aligns with our observations that increased synchronized TRN activity in *Msh2^-/-^* mice correlates with reduced sound evoked responses in the ACx and MGBv (Figure 1-3). Another factor that could contribute to the abnormal neural activity we observed in the TRN is increased parvalbumin expression, which has been shown to induce a change in firing patterns from regular to burst firing^65,66^. Consistent with this finding, burst firing in TRN neurons has been shown to be significantly altered in PV-KO mice compared to WT animals. In PV-KO mice, TRN neurons exhibited bursting epochs with prolonged ISIs compared to WT mice^25^. These observations complement our findings, as we observed that higher levels of PV in *Msh2^-/-^* mice correlated with shorter ISIs compared to *Msh2^+/+^* mice (Figure 6c).

Relatedly, the increase in burst firing epochs we observed in the TRN of *Msh2^-/-^* mice could also contribute to the PV^+^ interneuron swelling. Bursts of action potentials increase the influx of calcium and sodium ions into the intracellular space (Figure 5), potentially contributing to cellular edema^67^.

To delve deeper into the cellular pathologies associated with the TRN abnormalities, we analyzed acrolein levels, a marker of ROS and cellular oxidation that increases in the absence of DNA repair proteins^35^. Our analysis revealed significantly higher acrolein levels in the TRN of *Msh2^-/-^*compared to *Msh2^+/+^* (Figure 7). Since MSH2 is responsible for repairing base-base mismatches and insertion-deletion loops^4^, we speculate that its absence leads to increased DNA damage, which in turn raises ROS levels^68^. This increase in ROS leads to lipid oxidation, detectable by acrolein, and also results in oxidized DNA damage^69^. In *Msh2^-/-^* cells, the accumulation of oxidized DNA bases suggests that elevated oxidative stress is causing DNA damage that necessitates repair^9,36,37^. Without MSH2 to facilitate the repair of these oxidized bases, cells may activate neuroinflammatory pathways^70^, potentially impairing TRN function. Activation of neuroinflammatory pathways could have also contributed to the neuronal swelling observed in PV^+^ interneurons^71^. The intricate interplay of these factors creates a delicate balance within PV^+^ interneurons, which can quickly tip towards the development of pathological conditions.

In conclusion, our findings suggest that *Msh2* plays an important role in the maintenance of normal sensory processing. We have identified a potential involvement of *Msh2* in modulating the cortico-reticulo-thalamic circuit through the regulation of PV^+^ interneuron function. Future studies will elucidate the precise molecular mechanisms by which MMR promotes the development or maintenance of TRN neurons and provide valuable insights into how the brain utilizes DNA repair mechanisms for normal function.

## Supplementary Figure

**Figure S1:**
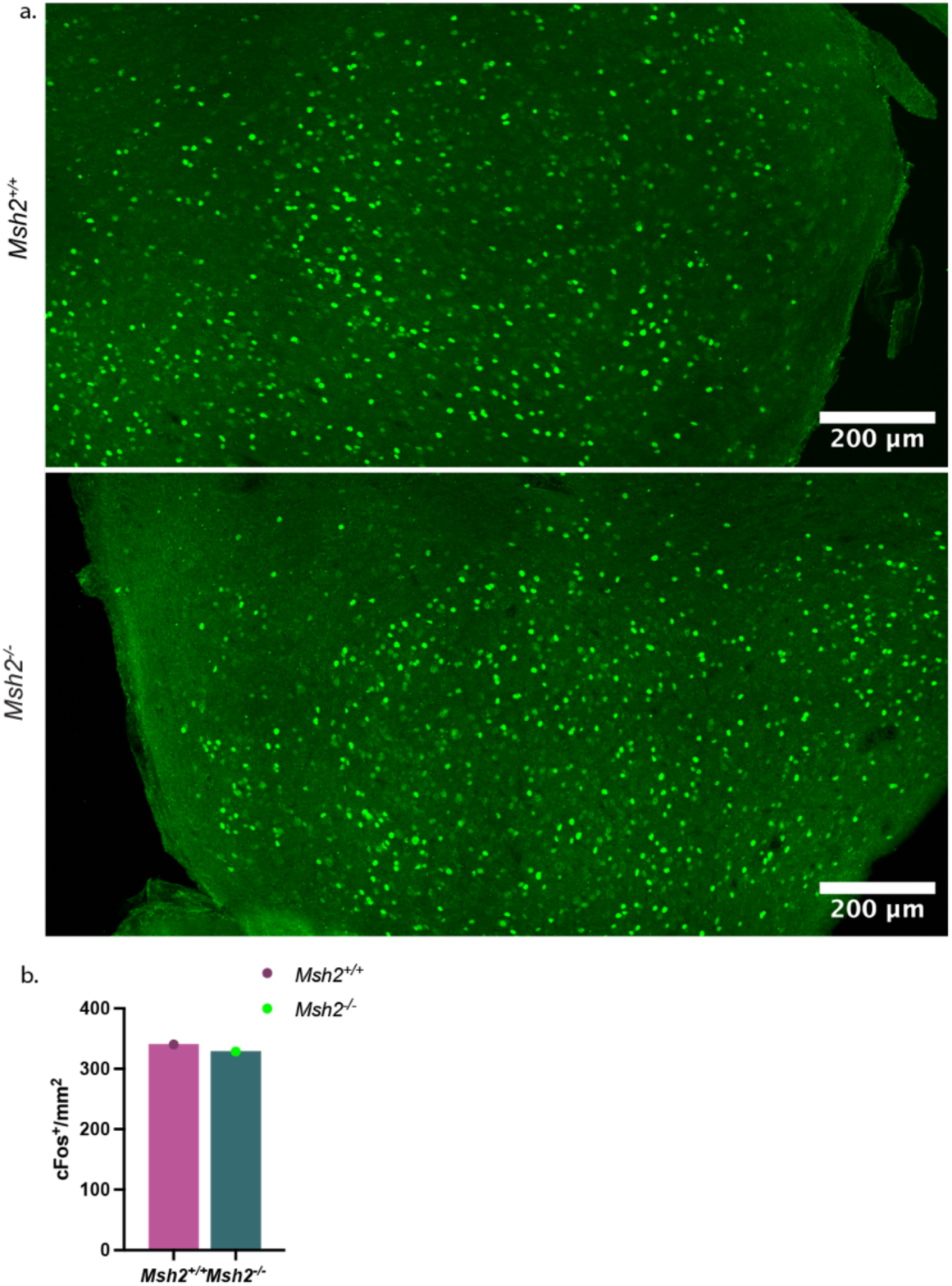
No difference in cfos expression in the inferior colliculus (IC) after auditory stimulation. **a)** cfos expression in *Msh2^+/+^* (top) and *Msh2^-/-^* (bottom) IC. Scale bars indicate 200µm. **b)** Quantification of cfos^+^ neurons in the IC (from data shown in panel a).

## Acknowledgements

We would like to thank Emily Sible, Mark Emerson and Pinar Ayata for valuable feedback on the project; Laura Nicolas-Gomariz for feedback on the manuscript. We would like to thank Gabriel Gray for troubleshooting the acrolein immunohistochemistry.

## Author contributions

S.N.R. performed and analyzed the cfos, PV, and connexin immunohistochemistry; D.N. performed and analyzed the in vivo electrophysiology experiments, S.O.G. performed and analyzed the acrolein immunohistochemistry; H.V.O. stimulated and perfused animals. S.N.R., H.V.O., and B.Q.V. drafted and edited the manuscript; B.Q.V. and H.V.O. coordinated the study.

